# The conserved ASCL1/MASH-1 ortholog HLH-3 specifies sex-specific ventral cord motor neuron fate in *C. elegans*

**DOI:** 10.1101/2020.06.04.134767

**Authors:** Lillian M. Perez, Aixa Alfonso

## Abstract

Neural specification can be regulated by one or many transcription factors. Here we identify a novel role for one conserved proneural factor, the bHLH protein HLH-3, implicated in the specification of sex-specific ventral cord motor neurons in *C. elegans*. In the process of characterizing the role of *hlh-3* in neural specification, we document that differentiation of the ventral cord type C neurons, VCs, within their motor neuron class, is dynamic in time and space. Expression of VC class-specific and subclass-specific identity genes is distinct through development and dependent on where they are along the A-P axis (and their position in proximity to the vulva). Our characterization of the expression of VC class and VC subclass-specific differentiation markers in the absence of *hlh-3* function reveals that VC fate specification, differentiation, and morphology requires *hlh-3* function. Finally, we conclude that *hlh-3* cell-autonomously specifies VC cell fate.

## INTRODUCTION

Cells in the nervous system, neurons and glia, are extremely diverse in shape, function, and the mechanisms by which they connect to other cells. Generation of neurons and their acquisition of unique features require the commitment to neural fate by an ectodermal descendant, the specification of neural class within the neuronal precursor, and the differentiation into unique transcriptomic and morphological states of the postmitotic cell. Importantly, the acquisition of pan-neuronal identity is seen to be regulated differently than the acquisition of unique neuronal class identity features. Redundant regulators with multiple cis-regulatory inputs induce pan-neuronal features whereas terminal differentiation of neurons is induced by single inputs, encoded by so-called terminal selectors, and results in the expression of a unique repertoire of genes that promote neural class diversity (Hobert, 2016b; Stefanakis et al., 2015). Thus, the neural diversity displayed by the nervous system is possible by the concerted action of terminal selector factors that function spatiotemporally with precision (Allan & Thor, 2015; Hobert, 2016a; Hobert & Kratsios, 2019; Kratsios et al., 2017). In *C. elegans*, the mechanisms that regulate neural specification can be studied thoroughly in time and in space, at single-cell resolution. This is a powerful model system that harbors a fully mapped body plan and nervous system, with continuously updated genomic and transcriptomic annotations, supporting studies in developmental biology, evolutionary conserved genes and networks, and beyond (Baker & Woollard, 2019; Cooper et al., 2018; Corsi et al., 2015; Emmons, 2016; Hammarlund et al., 2018; Sulston & Horvitz, 1997).

Here we characterize the role of a conserved proneural-like protein and ortholog of ASCL1/MASH-1, HLH-3, in *C. elegans* nervous system development. HLH-3 contains a conserved basic helix-loop-helix (bHLH) domain, which is 59% (31/54) identical to MASH1 and 61% identical to ASCL1 (33/54). HLH-3 heterodimerizes with the Class I bHLH transcription factor HLH-2, predicted ortholog of TCF3/TCF4/TCF12 (Kim et al., 2018; Krause et al., 1997). Our previous work has implicated HLH-3 in the terminal differentiation of the hermaphrodite-specific motor neurons, HSNs, a bilateral pair of neurons that function in the egg-laying circuitry (Doonan et al., 2008; Raut, 2017, Schafer, 2006). Work by others has shown that the gene *hlh-3* has diverse functions in the nervous system: it is necessary for the appropriate death of the sisters of the NSMs (Thellmann et al., 2003); it works in combination with other transcription factors to induce the serotonergic program in HSNs, and moreover, its ortholog, ASCL1, can be a functional substitute (Lloret-Fernández et al., 2018); it promotes neurogenesis of IL4 (Luo & Horvitz, 2017); it co-regulates the initiation of expression of the terminal selector gene *ttx-3* (Murgan et al., 2015); and it regulates the chemoreceptor gene *srh-234* (Gruner et al., 2016). Yet, one area that remains to be explored is the function of *hlh-3* in the post-embryonic ventral cord.

We were the first to report that *hlh-3* is expressed in the embryonically generated P cells, ectodermal-like precursors of all post-embryonically generated ventral cord motor neurons. We also showed that by the third larval stage (L3) expression of a truncated translational fusion *hlh-3* was restricted to the VCs, a hermaphrodite sex-specific type of neuron (Doonan et al., 2008). This expression pattern is consistent with a role in neuroblast specification, a function of canonical proneural proteins. However, it remained to be determined whether *hlh-3* had a function in the specification of their lineage descendants (including sex-shared as well as sex-specific). Here we report on the role of *hlh-3* in the development of these postembryonic ventral cord motor neurons. We show it is necessary for the acquisition and maturation of the hermaphrodite sex-specific VC class only.

The postembryonic ventral cord motor neurons are made up of both sex-specific and sex-shared neurons arising from the anterior descendants of ectodermal-like P blast cells (Pn.a) (Sulston & Horvitz, 1977). After two additional cell divisions, the Pn.aap cells give rise to the sex-specific neurons of the ventral cord. In hermaphrodites, the P3-P8.aap cells give rise to the ventral cord neuron type C (VC) (Figure 1A, B), whereas in males Pn.aapa and Pn.aapp (where n = descendant of P3 to P11), give rise to the ventral cord neuron type CA and CP, respectively (Sulston et al., 1980). Their fate acquisition (generation) is influenced by positional cues (Hox genes), differential survival (programmed cell death), and sexual identity (VC vs. CA/CP). The VCs of the hermaphrodite are positioned in the midbody and make up six of the total eight sex-specific neurons. Equivalent lineage descendants (Pn.aap) of P1, P2, and P9-12 cells in hermaphrodites undergo programmed cell death (Clark et al., 1993). Survival of VCs requires the function of the HOX gene *lin-39* and the HOX cofactors encoded by *unc-62* and *ceh-20* (Clark et al., 1993; Salser et al., 1993). UNC-62, along with LIN-39, promotes survival of the VCs by ensuring CEH-20 localizes to the nucleus; the LIN-39/CEH-20 complex then represses *egl-1* transcription (Liu et al., 2006; Potts et al., 2009). Sexual determination of the Pn.aap cells is established by the first larval (L1) stage (as VCs in hermaphrodites and the precursors of CAs and CPs in males) (Kalis et al., 2014). It was also shown that LIN-39 is not required for the expression of the VC terminal differentiation feature *ida-1*. Moreover, since the surviving descendants from P1, P2, and P9-12 still express *ida-1::gfp* in *lin-39 (lf); ced-3 (lf)* double mutants, it was concluded that the role of LIN-39 is most likely restricted to VC survival, not differentiation (Kalis et al., 2014). However, recent evidence has implicated a role for LIN-39 in the expression of a VC marker *srb-16* (Feng et al., 2020). Nevertheless, the mechanisms underlying VC class specification and differentiation are less understood. To date, it is not known which factor(s) initiate the differentiation program of VCs to establish a class-wide identity.

**Figure 1:**
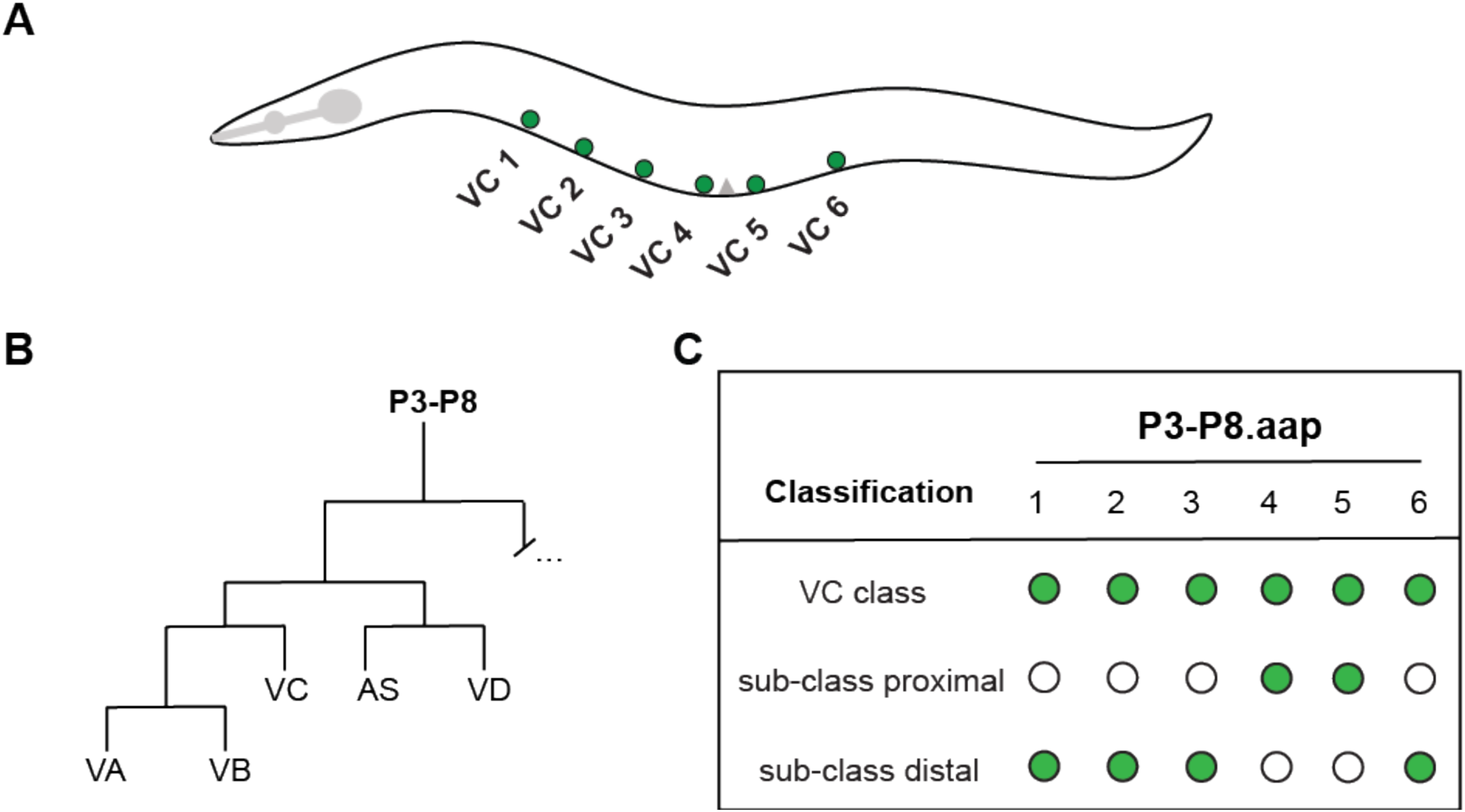
The ventral cord type C motor neuron class. **A:** Illustration of the position of the six VCs along the ventral nerve cord in the midbody region of an adult hermaphrodite. Anterior is to the left, ventral is down, gray triangle on the ventral surface indicates the location of the vulva. **B:** Diagram of the reiterative post-embryonic cell divisions produced by the P3.a to P8.a neuroblasts and give rise to the VCs (adapted from Sulston and Horvitz, 1977) **C:** Diagram for VC classification includes two sub-classes: proximal (VC 4 and VC 5) and distal (VCs 1-3, 6). This classification format will be used throughout the rest of the figures.

While the mechanisms that regulate VC class specification have yet to be determined, the mechanisms that regulate VC subclass identity are better understood. Within the VC class, two VC subclasses are distinguished spatially by their proximity to the vulva, categorized as proximal VCs or distal VCs (Schafer, 2006). The two VC neurons that flank the vulva are categorized as “proximal” (VC 4 and VC 5), whereas the other four VCs are “distal” to the vulva (VC 1-3, and VC 6) (Figure 1C). Genetic analysis of *unc-4* has revealed that VC subclass is determined by spatial cues.

Specifically, the expression of *unc-4* as a VC proximal subclass identity gene requires the secretion of EGF from vulval tissue (vulF cells) (Zheng et al., 2013). EGF signaling promotes proximal VC subclass fate by de-repression of *unc-4* in the proximal VCs only. Thus, a non-cell autonomous mechanism mediates one aspect of VC differentiation, specifically in proximal VCs.

Here we build on the current knowledge of neural specification in *C. elegans* and discover that the proneural-like bHLH factor, HLH-3, mediates specification and differentiation of the VC sex-specific motor neurons, that is, it is needed early and late in development. By using transcriptional reporter genes to assess VC differentiation in the absence of *hlh-3* function, we find that VC class and subclass identity, as well as morphology, is compromised. Our work is the first to identify a function for the ASCL1/MASH-1 ortholog, HLH-3, in the ventral cord, sex-specific neurons of *C. elegans* hermaphrodites. We conclude that HLH-3 is necessary for the expression of the earliest VC class-specific transcriptional regulator *(lin-11)* and is required for the expression of later acting VC class-specific genes.

## RESULTS

### The Class II bHLH protein HLH-3 is expressed and localized to the nuclei of VCs from L1 through adulthood

We have previously shown that in hermaphrodites, *hlh-3* is expressed in the postembryonic descendants of the ectodermal-like P cells as well as the HSNs (Doonan et al., 2008). We also have shown that *hlh-3* function is cell-autonomously required for normal axon pathfinding and terminal differentiation of the HSNs (Doonan, 2006; Doonan et al., 2008; Raut, 2017). In those studies, analysis of the expression of a translational fusion reporter with only the first eight amino acids of HLH-3 fused to GFP revealed that expression was widespread in the Pn.a descendants, dynamic, and with time, restricted to the VCs (Pn.aap) and HSNs. To confirm the endogenous spatiotemporal expression pattern of *hlh-3* we created *ic271 [hlh-3::gfp]*, a CRISPR-Cas-9 fluorescent tag at the C terminus of the *hlh-3* genomic locus (Figure 2A) following established genome-engineering protocols (Dickinson et al., 2015), and characterized its expression pattern. Our analysis supports our initial findings (Doonan, 2006; Doonan et al., 2008), the recently reported observation that *hlh-3* expression reappears in the HSNs at the L4 developmental stage (Lloret-Fernández et al., 2018), and expands our understanding of its role in the VCs (Doonan, 2006; Raut, 2017).

**Figure 2:**
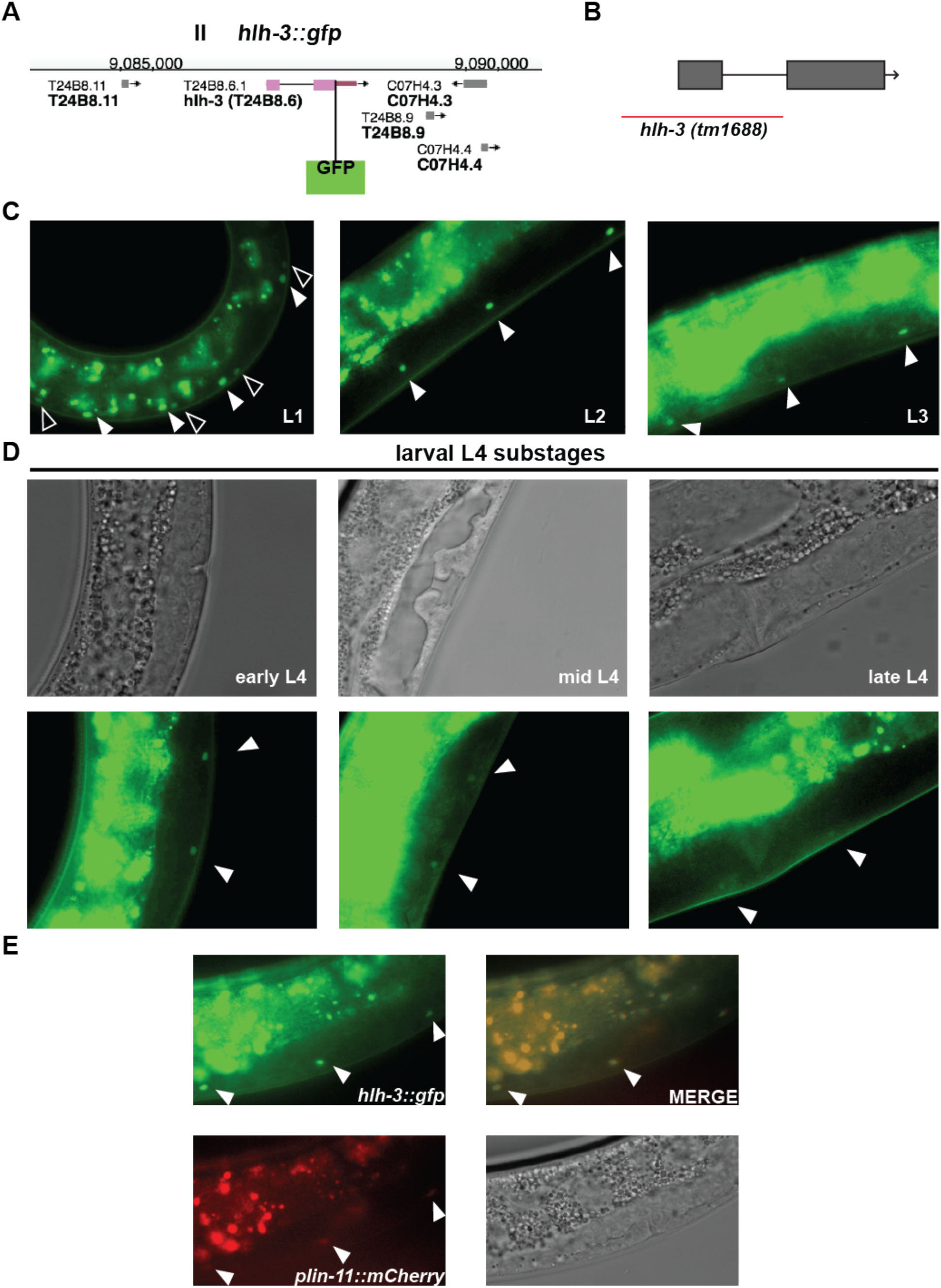
HLH-3 is first detected in nuclei of the Pn descendants (Pn.a and Pn.p) and becomes restricted to the nuclei of VCs as development proceeds. **A:** Diagram of the CRISPR-Cas9 engineered C-terminal GFP insertion at the *hlh-3* locus (*ic271*). **B:** The *hlh-3 (tm1688)* allele represents a 1242 bp deletion that spans chromosome II from 35,589 to 36,831 and removes exon 1. Removal of this region, including most of the bHLH domain, results in a null allele (Doonan et al., 2008). **C:** Representative images of the midbody ventral cord of hermaphrodites harboring HLH-3::GFP (*ic271*) at different larval developmental stages (L1, L2, and L3). At L1, filled arrowheads point to larger, more intense nuclei, presumably the Pn.a blast cells; whereas the outlined arrowheads point to the diminishing expression in Pn.p blast cells (left panel). Filled arrowheads in L2 and L3 represent expression in VC nuclei (middle and right panels). **D:** Representative images of the midbody ventral cord of hermaphrodites harboring HLH-3::GFP (*ic271*) over distinct L4 developmental stages (early, mid, and late). Larval substages (top panels) are classified by vulva morphology (Mok et al., 2015). Filled arrowheads point to the proximal VCs (bottom panels). **E:** Overlapping expression (merge, top right) of the VC marker, *plin-11::mCherry (otIs456)* (bottom left) and HLH-3::GFP *(ic271)* (top left), in an animal at the L4 molt (bottom right). Filled arrowheads point to co-labeled proximal VCs.

We confirmed that *hlh-3* is expressed post-embryonically in the P cells and their descendants and becomes restricted to the terminally differentiated VCs present in adults (Figure 2C, 2D, and 2E). After hatching, animals show the expression of *hlh-3* throughout the ventral nerve cord (VNC). We highlight the expression of *hlh-3* in an early L1 animal wherein Pn.p expression extinguishes faster than that in Pn.a and its descendants (Figure 2C, left panel). As development proceeds, expression is extinguished from other descendants of the Pn.a cells and restricted to the VCs (Figure 2C middle and right panels). While fluorescent reporter intensity was not quantified, *hlh-3* expression appears to be down-regulated in a window of the fourth larval stage (L4) development ranging from mid L4 to late L4, before increasing in adulthood (Figure 2D, middle and right panels). To ensure that the detected nuclei in adults are those of VCs, we characterized whether there was co-expression of *hlh-3::gfp* with *plin-11::mCherry*, a known VC marker (Figure 2E, bottom left). We find that the *hlh-3::gfp* positive nuclei are also *plin-11::mCherry* positive (Figure 2E, top right panel). Interestingly, low levels of *hlh-3::gfp* expression is also observed in a pair of vulval cells during mid-late substages of L4 development, suggesting a role for *hlh-3* in these lineages (data not shown). Expression of *hlh-3* in VCs from their birth in L1 through their terminally differentiated stage in adulthood prompted us to investigate the role of *hlh-3*, as a factor required for an early role in promoting VC fate and required for maintenance of VC fate throughout development. Throughout we will use the allele *hlh-3 (tm1688)*, which eliminates the majority of the bHLH domain and transcription start site rendering this a null allele and further referred to in this paper as *hlh-3 (lf)* (Figure 2B, Doonan et al., 2008).

### Differentiation of VC class and VC subclass motor neurons is dynamic

Before we analyzed the role *hlh-3* in VCs, we first characterized differentiation features of VCs by defining VC class versus VC subclass-specific terminal identity. Here we took advantage of fluorescent reporter genes that serve as markers of VC fate. We examined the expression of the known VC class-specific markers *lin-11, ida-1*, and *glr-5* encoding a LIM homeodomain transcription factor (Freyd et al., 1990), a protein tyrosine phosphatase-like receptor protein homolog of IA2 (Cai et al., 2001, 2004; Zahn et al., 2001), and a glutamate receptor subunit (Brockie et al., 2001), respectively. We confirmed that expression of *lin-11* in VCs is observed as early as the second larval stage (L2), and through adulthood (data not shown) (Hobert et al., 1998; Zheng et al., 2013).

Unlike *lin-11*, a transcriptional regulator, the other VC class terminal identity genes *ida-1* and *glr-5* are expressed later in development, arising at the L4 developmental stage (Figure 3A and 3B). Analysis of these VC class differentiation markers throughout L4 substages revealed distinct spatiotemporal patterns suggesting different pathways regulate them. Expression of *ida-1* and *glr-5* is not equivalent across all 6 VCs during L4 development (Figure 3A and 3B). Classification of the L4 substages (early, mid, and late) is based on the vulval L4 morphology as previously described (Mok et al., 2015). We noted that *glr-5* expression is first detected in the early L4 substages and only in the proximal VC subclass, whereas expression can be detected in the distal VCs by late L4 substages (Figure 3B). In contrast to *glr-5, ida-1* expression is nearly equivalent in all VCs since the beginning of L4, but its expression is always detectable in the posterior VCs (Figure 3A). Thus, while the six VCs terminally express their class-specific terminal differentiation genes *ida-1* and *glr-5*, the initiation of transcription is distinct across the sub-stages of L4 development.

**Figure 3.**
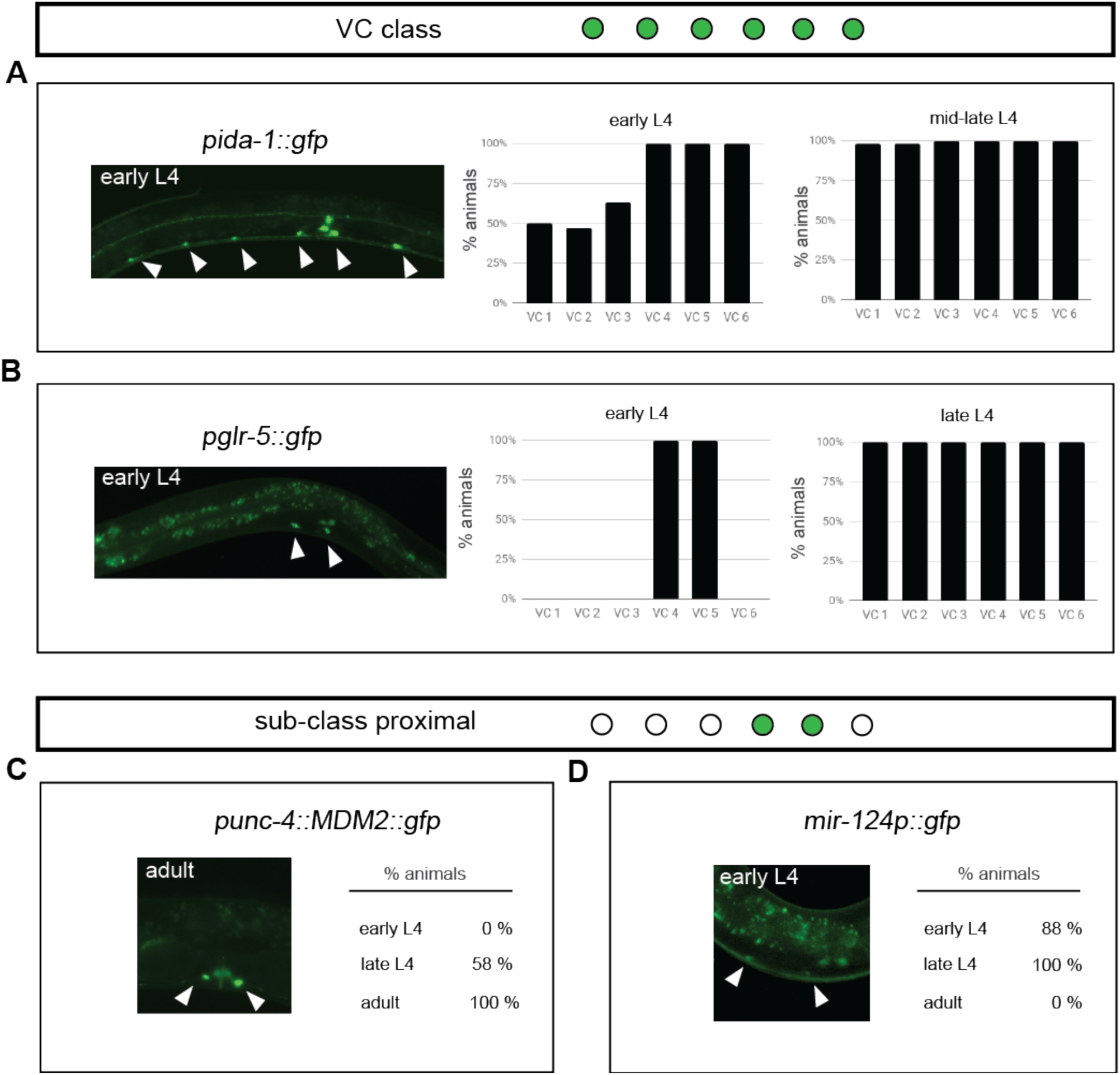
The spatiotemporal expression of VC class and subclass-specific identity features is dynamic. **A:** Expression pattern of the VC class differentiation feature *ida-1* in early and mid-late L4 developmental stages. Image shows an early L4 hermaphrodite expressing *inIs179 [pida-1::gfp]* in all VCs (indicated by arrowheads). Graphs report the percent of animals expressing *inIs179 [pida-1::gfp]* (early L4, n = 20; mid-late L4, n = 40) in each VC. Since all VCs express *pida-1::gfp* by mid-L4, the sub-stages in late L4 were grouped together with mid-L4 substages. **B:** Expression pattern of the VC class differentiation feature *glr-5* in early and late L4 development. Image shows an early L4 hermaphrodite expressing *icIs270 [pglr-5::gfp]* in the proximal VCs (indicated by arrowheads). Graphs report the percent of animals expressing *icIs270* in early L4 (n = 10), mid L4 (n = 15), and late L4 (n = 15) developmental stages in each VC. **C:** Quantification of expression of the VC subclass feature, *unc-4*, from L4 development through adulthood. Image shows the expression of *uIs45 [punc-4::MDM2::GFP]* in an adult (indicated by arrowheads). The percent of animals expressing the VC subclass marker in both cells (VC 4 and VC 5) of early L4 (n = 8), late L4 (n = 12), and adults (n = 19). **D:** Quantification of expression of VC subclass feature, *mir-124*, from L4 development to adulthood. Image shows expression of *mjIs27 [mir-124p::gfp + lin-15(+)]* in the proximal VCs of an early L4 hermaphrodite. Percent of animals expressing the VC subclass marker in both cells during these substages is listed adjacent to the image in early L4 (n = 8), late L4 (n = 13), and adults (n = 10).

Next, we characterized the expression pattern of the VC subclass-specific terminal identity genes *unc-4* and *unc-17*. Others have shown that *unc-4* expression requires *lin-11* and vulval EGF signaling (Zheng et al., 2013). We corroborate that *unc-4* expression is detected after the mid-L4 stages and is maintained throughout adulthood only in VC 4 and VC 5 (Figure 3C). The expression of UNC-17, in turn, is known to require a posttranscriptional step mediated by UNC-4 (Lickteig et al., 2001). Therefore, we analyzed the expression of two transcriptional *unc-17* reporters. To our surprise, and in contrast to work by others, we only detect the expression of *unc-17* in VC 4 and VC 5 at the adult stage regardless of which reporter we characterized (Supplemental Figure 1) (Pereira et al., 2015). However, our work is different from others in that we did not assess a translational reporter. Instead, we looked at two transcriptional reporters *vsIs48 (punc-17::gfp)* and *mdEx865 (unc-17p::NLS::mCherry + pha-1(+))* and did not observe *unc-17* expression in the distal VCs 1-3 and 6 with either reporter (*vsIs48* expression is shown in Supplemental Figure 1B top panel; *mdEx865* expression is not shown). Although we do not see the *unc-17* reporters in the distal VCs we still detect a VC marker (*lin-11*) in these cells (Supplementary Figure 1B middle and bottom panels). Our observations are also consistent with previous reports that anti-UNC-17 immunoreactivity is robust in VC 4 and VC 5, but rarely detectable in distal VCs (Duerr et al., 2008; Lickteig et al., 2001) and possibly only in the second larval (L2) stage (Alfonso et al., 1993).

### *mir-124* is a novel VC subclass-specific identity feature

In our search for VC subclass identity genes, we found *mir-124*, the highly conserved non-coding microRNA, as a novel VC subclass-specific differentiation feature. In *C. elegans* it has been documented to be expressed in a variety of sensory neurons and the HSNs (Clark et al., 2010). Here we characterized *mir-124* expression across postembryonic development; we only see it in a restricted window. We find *mir-124* is expressed from early L4 larval substages through early adulthood, but not in mature gravid egg-laying hermaphrodites (Figure 3D), which suggests it is required for the maturation of the VCs but not for maintenance of VC fate. This expression pattern is unlike that of other proximal VC identity features *unc-4* and *unc-17*, which are expressed throughout adulthood (Figure 3C, Supplemental Figure 1). Therefore, we classify *mir-124* as a novel VC subclass-specific feature expressed during early differentiation. In summary, we conclude that *mir-124* can be added to the list of VC identity features belonging to the proximal class (Figure 4).

**Figure 4:**
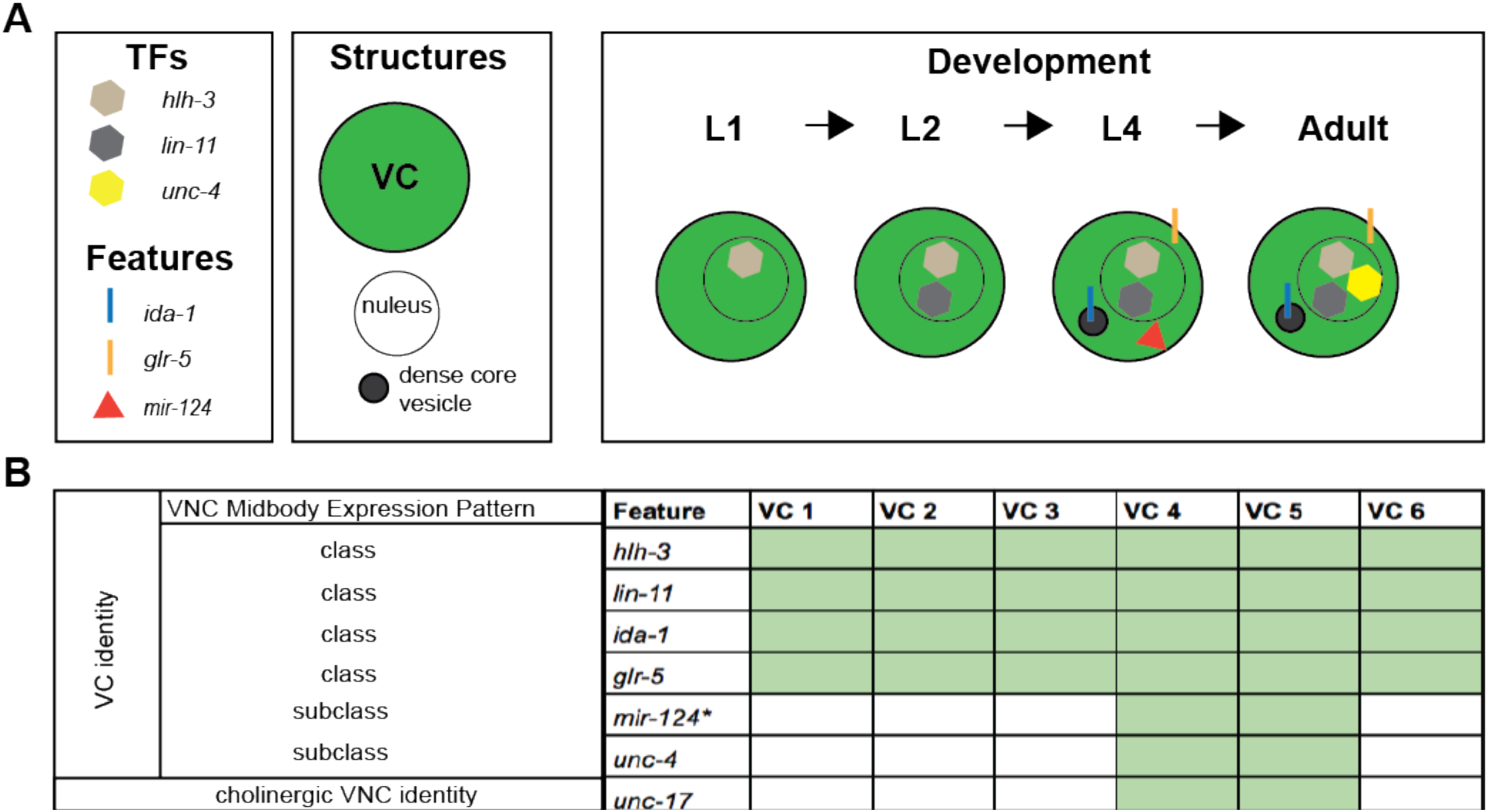
Summary of VC class and subclass identity. **A:** Diagram of genes encoding transcription factors (TFs) and class- or subclass-specific features and structures expressed in VCs throughout post-embryonic development. **B:** Summary of the expression pattern of distinct VC class and VC subclass identity features in the midbody of the ventral cord of the hermaphrodite. While *unc-17* is expressed in the subclass proximal VCs, it is also expressed in all VNC cholinergic motor neurons, therefore not VC specific. Our analysis is based on the expression of integrated transcriptional reporters with the exception of the endogenous GFP tag to *hlh-3* (See Supplemental Table 1 for the list of strains containing these markers). With the exception of *mir-124*, all reported features are maintained through adulthood.

### Classification of VC identity

Thus far, we have shown that the VC class of neurons acquire class-specific features via mechanisms that differ in time and space. We have also shown that not all six VCs are identical in their repertoire of transcriptional activity. In Figure 4, we summarize the spatiotemporal expression pattern of VC identity features. Two genes encoding presumptive transcription factors, *hlh-3* and *lin-11*, are VC class-specific and *hlh-3* expression precedes that of *lin-11* (Figure 4A-C). We classify the *ida-1* and *glr-5* genes as VC class-specific as well, as they are observed in all six VCs from L4 through adulthood. In contrast, *mir-124* is not expressed in adulthood, as the rest of the VC terminal identity features are. Finally, *unc-17* was observed in the proximal VCs only. It is worth emphasizing that aside from the analysis of the CRISPR-engineered *hlh-3::gfp* line, our analysis is based on the characterization of transcriptional reporters (Supplementary Table 1).

### *hlh-3* function is required for the acquisition of VC class and VC subclass identity features

Previously, *hlh-3* has been shown to be required for HSN terminal differentiation (Doonan et al., 2008; Lloret-Fernández et al., 2018; Raut, 2017). To address whether *hlh-3* has a role in VC differentiation we first examined reporters of VC terminal identity the genes *lin-11, ida-1* and *glr-5*, in a strain harboring a total loss of *hlh-3* function allele, *hlh-3 (tm1688)* (Doonan et al., 2008, Figure 2B). We find that expression of the terminal VC class markers *lin-11, ida-1* and *glr-5* is reduced in most of the VCs of one day old *hlh-3 (lf)* adult hermaphrodites (Figure 5A, 5B, and 5C). In fact, expression of the differentiation features *ida-1* and *glr-5* (not shown) in the earlier stages of L4 development is completely absent in *hlh-3 (lf)* hermaphrodites (Supplementary Figure 2 A and B).

**Figure 5.**
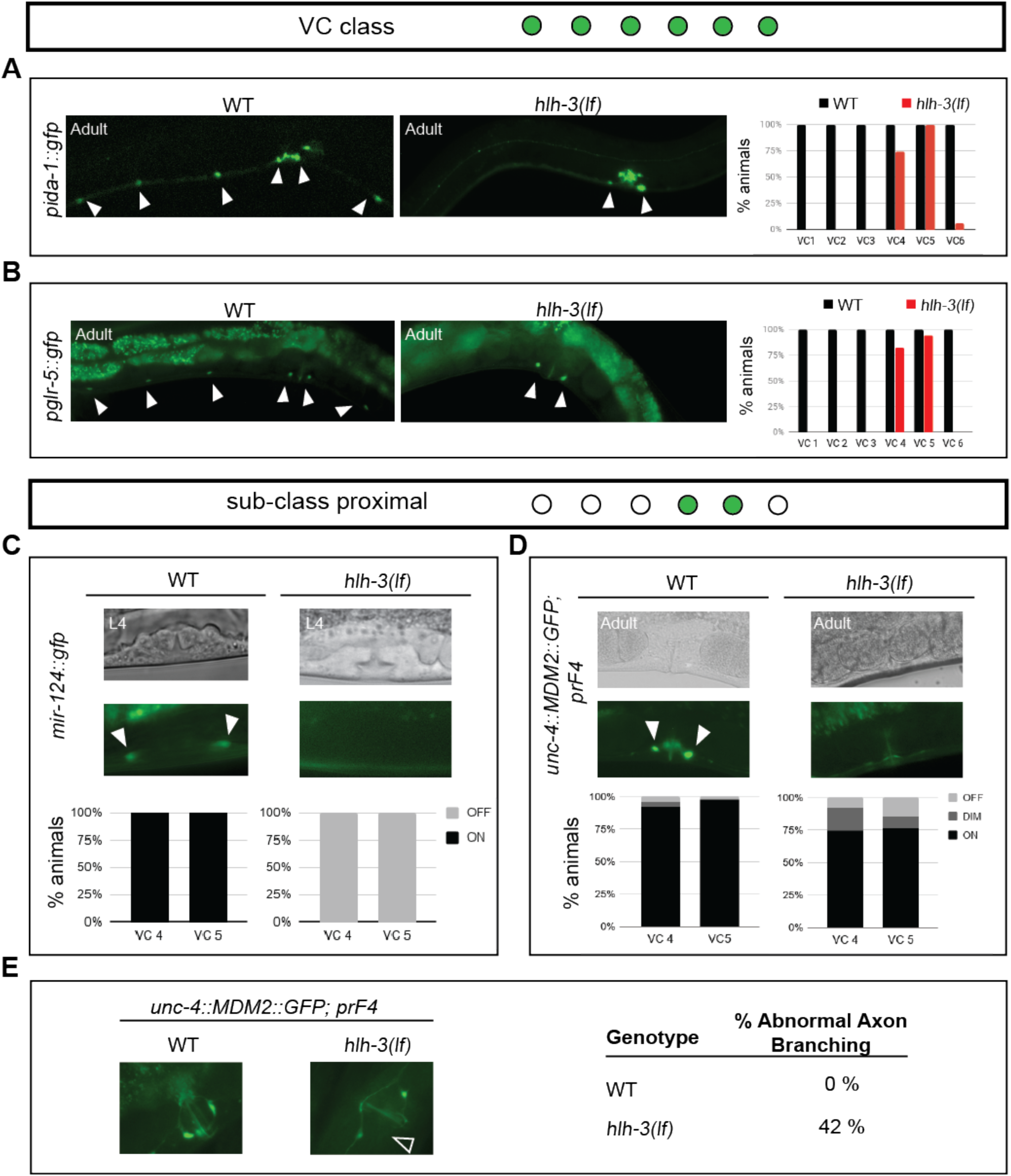
VCs require HLH-3 to acquire class-specific and subclass-specific differentiation features and normal axon morphology. **A-B:** Representative images of WT or *hlh-3 (lf)* individuals harboring the indicated reporters. Filled arrowheads point to detectable VCs in either genotype. Graphs report the percent of animals expressing each reporter in each VC of WT (gray bars) and *hlh-3 (lf)* (red bars). The expression of the *ida-1* marker (*inIs179*) was quantified in WT (n = 15) and *hlh-3 (lf)* (n = 35) (panel A). The expression of the *glr-5* marker (*icIs270*) was quantified in WT (n = 15) and *hlh-3 (lf)* (n = 30) (panel B). **C-D:** Representative images of WT and mutant hermaphrodites at different stages of development and harboring the indicated reporters of VC subclass features *mir-124 (mjIs27)* and *unc-4 (uIs45)*. Fluorescent images of the vulval region of DIC imaged hermaphrodites (top panels) only revealed expression in the proximal VCs of WT individuals (indicated by filled arrowheads). Quantification of the percent of animals with detectable reporter expression of *mir-124* (*mjIs27*) in VC 4 or VC 5 is reported in the graph below the images. Fluorescence was either detectable (on) or not detectable (off) for expression of *mjIs27* in WT mid L4s (n = 17) and *hlh-3 (lf)* mid L4s (n = 14). Quantification of the percent of animals with detectable reporter expression of *uIs45* in VC 4 or VC 5 is reported in the graph below the images. Fluorescence was either bright (on), dim, or not detectable (off) for expression of *uIs45* in WT (n = 66) and *hlh-3 (lf)* (n = 61) adults. **E:** Quantification of proximal VC axon branching in WT and *hlh-3 (lf)* individuals. Normal axons branch into a vulval ring, as observed with *uIs45* in the WT genotype (top panel). In contrast, *hlh-3 (lf)* hermaphrodites display abnormal axon branching (bottom panel). The numbers to the right represent the percent of individuals with abnormal branching in adult WT (n = 15) and *hlh-3(lf)* (n = 24) adult hermaphrodites.

Next, we examined the expression of VC subclass-specific identity features, *mir-124, unc-4*, and *unc-17* (Figure 5 and Supplemental Figure 1). We find that the early differentiation subclass-specific feature *mir-124 (mjIs27: mir-124p::gfp + lin-15(+))* is completely absent in *hlh-3 (lf)* (Figure 5D). We followed up with an analysis of *unc-4*. Others have shown that the expression of this VC subclass-specific terminal identity gene is de-repressed in WT animals after EGF signaling in mid-L4 development (Zheng et al., 2013). Here, we find that the absence of *hlh-3* function negatively affects *unc-4* expression (Figure 5E). Since *unc-4* expression is required for *unc-17* expression (Lickteig et al., 2001), not surprisingly we find that expression of *unc-17*, is missing the proximal VCs in *hlh-3 (lf)* individuals (Supplemental Figure 1E).

### *hlh-3* is required for normal axon branching of proximal VCs

Along with less expression of VC terminal identity transcriptional reporters, proximal VCs have abnormal axonal branching in the vulval ring (Figure 5F). This defect suggests that proximal VC function may be impaired in *hlh-3 (lf)*, as axonal branching is required for synaptic connections to the egg-laying circuitry. Thus, growth and maturation of VC axons require *hlh-3* function, as it is the case for the HSNs (Doonan, 2006; Doonan et al., 2008; Raut, 2017).

### VCs survive in the absence of *hlh-3* function

Our analysis of VC class and VC subclass markers indicate that the expression of VC differentiation markers is compromised in *hlh-3 (lf)* individuals. To ensure VC survival occurs we next sought to eliminate the possibility that VCs inappropriately undergo programmed cell death in *hlh-3 (lf)*. Programmed cell death (PCD) is a conserved pathway executed by CED-3, a caspase that functions as the final determinant in the cell death pathway (Conradt et al., 2016). Inhibition of this pathway, by impairment of *ced-3* function, results in the survival of cells destined to die. In the context of the ventral nerve cord, the cells P1-P2.aap and P9-12.aap will survive (Figure 6A). Therefore, we introduced a *ced-3* null mutation into *hlh-3 (lf)* mutants and analyzed the expression of a VC differentiation marker, *grl-5*, in *ced-3 (lf)* and *ced-3 (lf); hlh-3 (lf)* individuals. Unlike *ced-3 (lf)* hermaphrodites, which express *glr-5* in all VCs including the surviving P2.aap cell, we find that *ced-3 (lf); hlh-3 (lf)* mutants do not express *glr-5* in VCs or the surviving VC-like cell P2.aap (Figure 6B, 6C, and 6D). Therefore, we conclude that the reason VC neurons do not express *glr-5* in the absence of *hlh-3* function is that they need HLH-3 to fully differentiate and not because they undergo inappropriate PCD.

**Figure 6.**
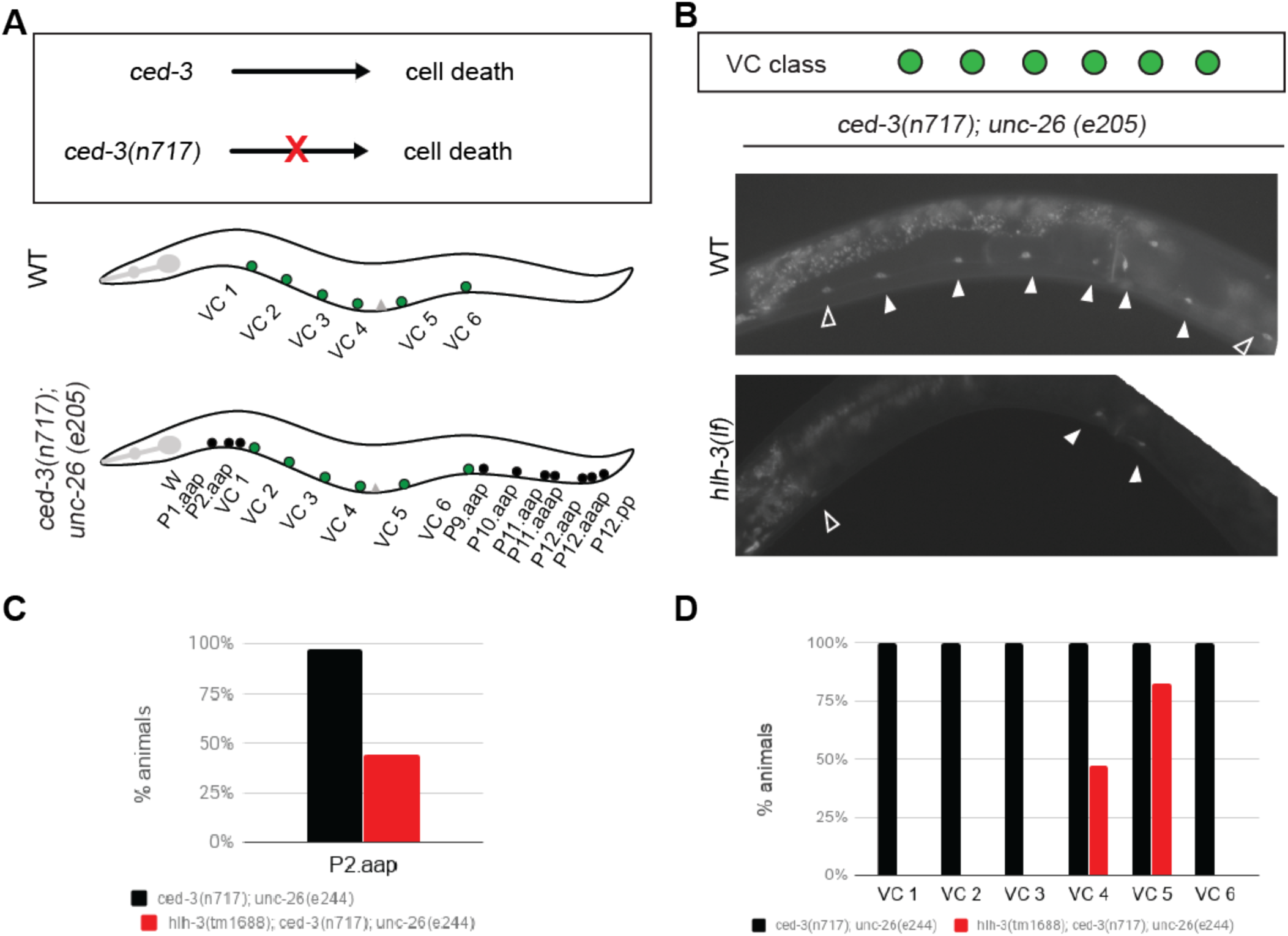
VCs do not inappropriately undergo programmed cell death (PCD) in the absence of *hlh-3* function. **A:** Diagram of outcome in the presence and absence of *ced-3* function. The presence of *ced-3* function in WT individuals results in PCD, the absence of *ced-3* function in the null allele *ced-3* (*n717*) prevents PCD. In the ventral nerve cord of WT animals, only the descendants of P3.aap to P8.aap or VCs report the expression of VC markers. However, in *ced-3 (n717)* nulls, the VC-equivalent descendants of P1 and P9-12, that normally undergo PCD, do not undergo PCD and report expression of VC markers. **B:** Representative images of *ced-3 (n717); unc-26 (e244)* individuals with (WT), or without (*hlh-3 (lf))* function. The *glr-5* VC marker (*icIs270*) was utilized to monitor the presence of VCs (filled arrowheads) and VC-like surviving cells (outlined arrowheads; specifically, P2.aap and P9.aap). The reporter *icIs270* is only detected in the proximal VCs (filled arrowheads) and a VC-like cell (outlined arrowhead; P2.aap) in the double mutant *hlh-3 (lf)*; *ced-3 (lf)*. **C:** Quantification of the percent of one day old adults expressing *icIs270* in P2.aap in *ced-3 (n717); unc-26 (e244); pglr-5::gfp* (n = 35), and *hlh-3 (tm1668); ced-3 (n717); unc-26 (e244); pglr-5::gfp* (n = 34). **D:** Quantification of the percent of one day old adults expressing *icIs270* in each VC of *ced-3 (n717); unc-26 (e244); pglr-5::gfp* (n = 35) and *hlh-3 (tm1668); ced-3 (n717); unc-26 (e244); pglr-5::gfp* (n = 34).

### *hlh-3* functions cell-autonomously in the VC class

To address whether *hlh-3* functions cell-autonomously, we assayed expression of a VC differentiation marker *plin-11::mCherry* in *hlh-3 (lf)* mutants with a rescuing copy of *hlh-3*. The rescuing extrachromosomal array *[icEx274 (plin-11::pes-10::hlh-3cDNA::GFP; pmyo-2::mCherry)]* was made by introducing a *hlh-3* cDNA into pDM4 (previously shared by Michael Koelle) harboring a VC-specific regulatory region of *lin-11* fused to the basal *pes-10* promoter (Doonan, 2006). We find that whereas *hlh-3 (lf)* mutants fail to express the VC differentiation marker *plin-11::mCherry* in most VCs, *hlh-3 (lf)* mutants that contain the rescuing extrachromosomal array *icEx27*4 express *plin-11::mCherry* in almost all VCs (Figure 7A and B). These findings demonstrate that *hlh-3* function is cell-autonomous.

**Figure 7.**
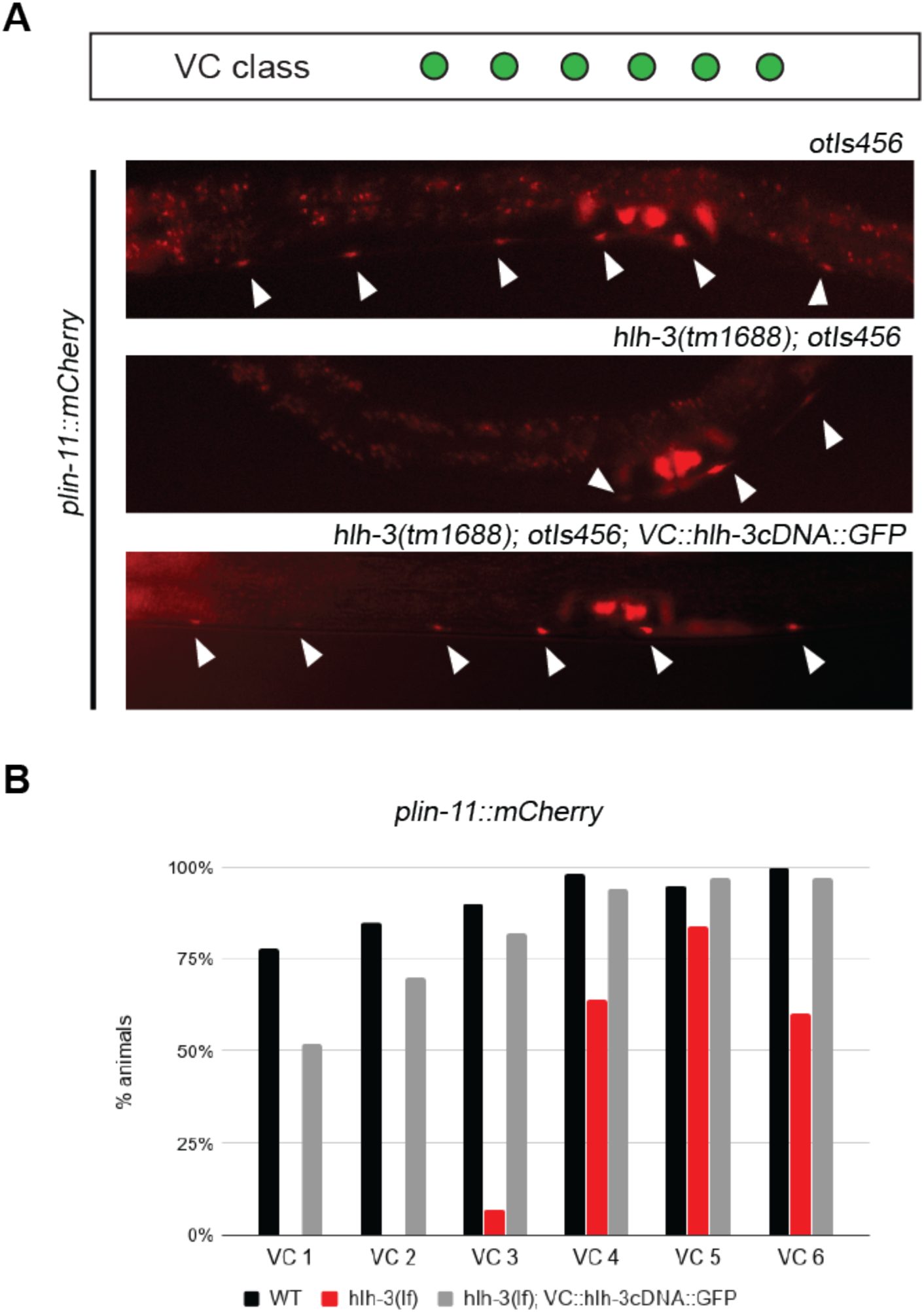
The function of *hlh-3* in VCs is cell-autonomous. **A:** Representative images of individuals harboring the *lin-11* marker (*plin-11::mCherry*) in WT (top panel), *hlh-3 (lf)* (middle panel), and VC-specific rescued lines (bottom panel). The reporter *otIs456* is normally expressed in all VCs (top panel, filled arrowheads). **B:** Quantification of the percent of mid-late L4 animals expressing *plin-11::mCherry* in each VC of WT (n = 41), *hlh-3 (lf)* (n = 48), and *hlh-3 (lf); VC::hlh-3cDNA::GFP* (n = 39).

### *hlh-3* does not affect the differentiation of other sex-shared neurons in the ventral cord

Given that expression of the *hlh-3* CRISPR-edited reporter is detectable in the P cells and its descendants we wished to address whether the absence of *hlh-3* function resulted in defects in the sex-shared neurons. To address this question, we analyzed the expression of cholinergic and GABAergic markers in *hlh-3 (lf)* mutant hermaphrodites. The transcriptional reporter *vsIs48 [punc-17::gfp]* gene marks all cholinergic neurons expressing a vesicular acetylcholine transporter (within the VNC this includes VA, VB, AS, DA, DB, Supplemental Figure 1A) (Wormbase: Curatorial remark). The transcriptional reporter *otIs564 [punc-47::mChOpti]* marks all GABAergic neurons expressing a vesicular GABA transporter (within the VNC this includes DD and VD neurons, Supplemental Figure 3A) (Gendrel et al., 2016). We find that the total number of cholinergic neurons anterior to the vulva is equivalent between WT and *hlh-3 (lf)* individuals (Supplemental Figure 1C). Likewise, the total number of GABAergic neurons is equivalent between WT and *hlh-3 (lf)* hermaphrodites (Supplemental Figure 3B). These analyses demonstrate that the cholinergic and GABAergic sex-shared ventral cord motor neurons acquire their terminal neurotransmitter fate. Thus, *hlh-3* function is not necessary for the acquisition of the terminal fates in sex-shared neurons, rendering its function specific to the terminal differentiation of sex-specific ventral cord VC neurons.

### The male-specific ventral cord motor neurons do not require *hlh-3* function

We wondered whether the male-specific ventral cord motor neuron differentiation was also dependent on *hlh-3* function. The CA and CP pairs of male motor neurons arise from the division of the Pn.aap neuroblast, anteriorly (type CA) and posteriorly (type CP) (Sulston et al., 1980; Supplementary Figure 4A). We tracked differentiation of the Cas 1-9 and the CPs 1-6 with the differentiation markers for *ida-1 and tph-1*, respectively (Supplementary Figure 4B). We find that the *hlh-3 (lf)* males when compared to WT males show expression of differentiation markers in all CA and CP neurons, nearly at equivalent proportions (Supplementary Figure 4C and 4D). This suggests that *hlh-3* does not have a role in promoting the differentiation of these neurons. Notably, we did not quantify CP0 (descendant of the P2 lineage) although we did observe expression of *pida-1::gfp* in both WT males and *hlh-3 (lf)* (data not shown).

## DISCUSSION

### *hlh-3* specifies VC fate

Our work identifies *hlh-3* as a regulator of sex-specific motor neuron differentiation in the postembryonic VNC of the hermaphrodite. Both terminal and non-terminal identity features associated with the sex-specific motor neurons, VCs, are reduced or absent in animals that lack *hlh-3* function. While most of our analysis measures transcriptional gene activity of VC identity genes, we also demonstrate that the morphology of the VC subclass is affected. In summary, we implicate *hlh-3* in the specification of the VC motor neuron class.

### Differentiation of the proximal VCs involve *hlh-3* dependent and *hlh-3*-independent mechanisms

Our work demonstrates that in the absence of *hlh-3* function, the differentiation of proximal VCs is less affected than that of distal VCs. We have gained some insight into these differences with the analysis of markers that are expressed in early L4 versus later L4 substages (Figure 3). Expression of VC class and VC subclass-specific identity features *ida-1, glr-5*, and *mir-124*, is seen in VCs in early L4 substages in a WT context, yet, are completely absent from these early substages through adulthood in animals that lack *hlh-3* function (Figure 3, Figure 5, Figure 8A, Supplemental Figure 2). This indicates *hlh-3* function is required before L4 development. We also learned that in the mutant context, and during later stages of L4 development, expression of these VC identity features appeared in just a few VCs, the proximal ones. This suggests that there may be a parallel pathway, which can promote VC differentiation. Since the proximal VCs are less affected in their expression of the terminal identity genes that arise after mid-L4 development (*unc-4* and *unc-17*), we propose that this alternative pathway acts by mid L4 but not sooner. We infer that the *hlh-3* independent parallel pathway is mediated by EGF, a cue secreted as early as mid L4, already shown to be required for expression of *unc-4* in proximal VCs (Figure 8B; Zheng et al., 2013). The presence of this parallel pathway could ensure that at least proximal VCs retain some function, as they are primary contributors to egg-laying by providing feedback to HSNs and vulva muscles (Schafer, 2006).

**Figure 8.**
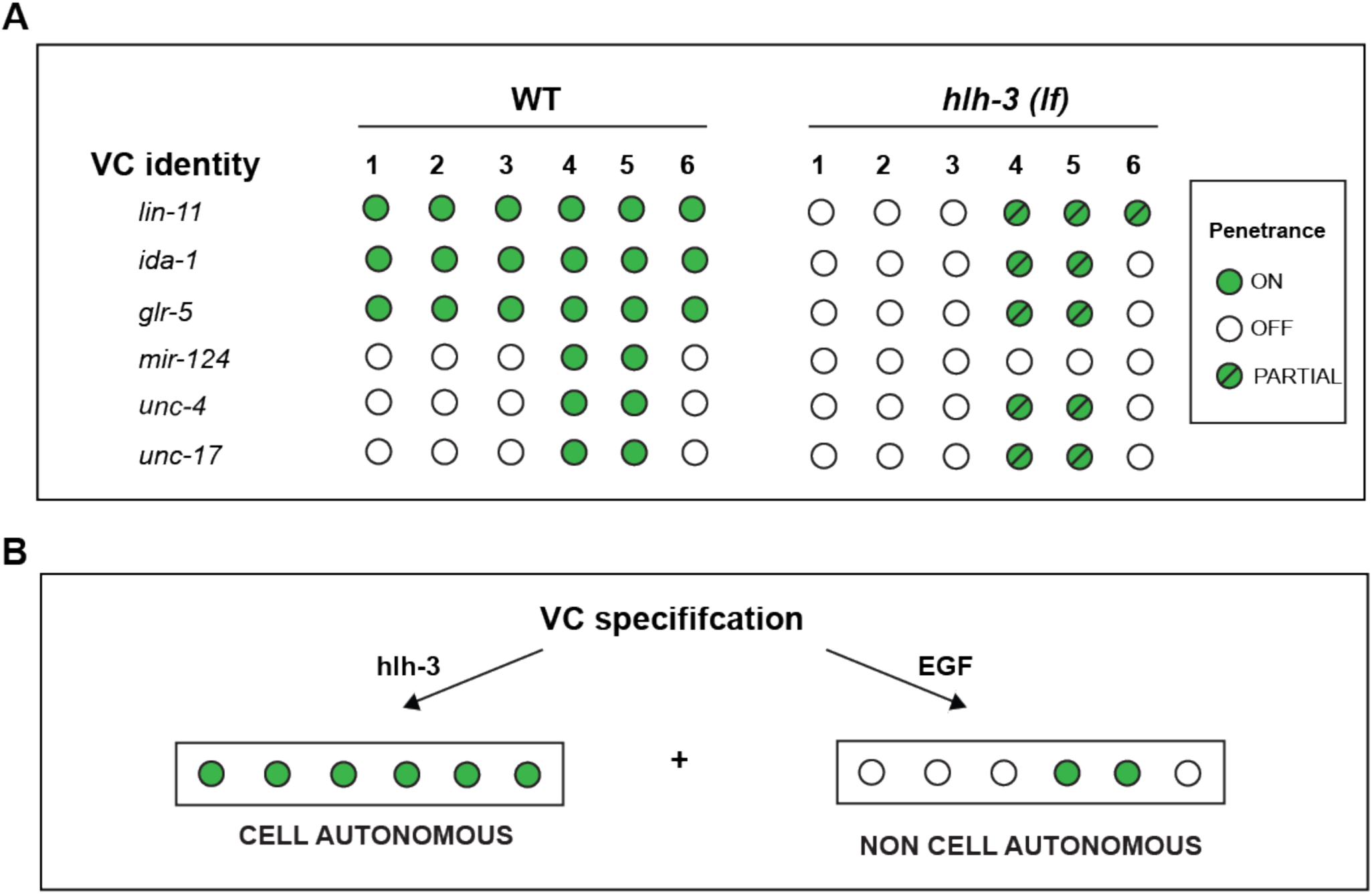
Two pathways promote the acquisition and maintenance of VC-class and VC-subclass features. **A:** Expression of the VC identity features (*lin-11, ida-1, glr-5, mir-124*) require *hlh-3* function. **B:** The regulation of the VC identity features occurs in a cell-autonomous way prior and independently of EGF signaling during mid-L4 development. The alternative pathway, dependent on EGF, regulates the expression of *unc-4* (Zheng et al., 2013). We propose that the function of EGF signaling adds a secondary input to regulate *lin-11* levels in the proximal VCs, and affect *unc-4* and *unc-17*, as well as other VC identity features.

In summary, we have found that the acquisition of VC class features (shown herein) is impaired in *hlh-3 (lf)* individuals. Of the features we have analyzed, only one subclass differentiation feature, expression of *mir-124*, is fully dependent on *hlh-3* function (Figure 5C, Figure 8A). Since *mir-124* expression is restricted to the VC proximal subclass, it may have a role in promoting VC subclass diversity. However, since the expression of *mir-124*, is seen prior to the EGF cue, and is completely absent in *hlh-3 (lf)*, we believe that it is regulated by *hlh-3* and not by the EGF-dependent pathway. Further work has to address whether *mir-124* functions as an intrinsic, cell-autonomous mechanism to promote VC class diversity.

With this work, we propose that: (1) *hlh-3* functions cell-autonomously to specify VC class fate early in development (from L1 to L4) and (2) during L4 development an EGF-dependent cue promotes proximal VC subclass fate diversity for function in egg-laying. Our proposal is consistent with the observation that expression of *lin-11, glr-5, ida-1*, and *unc-4*, in the proximal VCs, is not significantly altered in the absence of *hlh-3* function. To reiterate, the proposed *hlh-3* dependent pathway specifies VC class fate and an *hlh-3* independent pathway promotes VC subclass diversity.

### The LIM homeodomain transcription factor LIN-11 in VCs is downstream of and positively regulated by *hlh-3*

As shown by others, the gene encoding LIN-11 is expressed from L2 through adulthood (Hobert et al., 1998). We have observed this as well with the translational reporter *wgIs62 (lin-11::TY1::EGFP::3xFLAG + unc-119(+))* (data not shown). Since our analysis indicates that *hlh-3* is expressed before *lin-11*, we characterized the expression of a *lin-11* transcriptional reporter *(plin-11::mCherry)* in the absence of *hlh-3* function. We showed that *hlh-3 (lf)* mutants exhibit reduced *lin-11* transcriptional activity in VCs (Figure 7). It is likely that *hlh-3* directly targets *lin-11*, but further work will determine whether this effect is indirect or indirect. Interestingly, the ortholog ASCL1 has been shown to directly target the *lin-11* ortholog, Lhx1, in a ChIP-seq analysis of the ventral telencephalon (Castro et al., 2011; Kim et al., 2018).

Our analysis of *lin-11* expression in *hlh-3 (lf)* also revealed that the proximal VCs are less affected than the distal VCs by the absence of *hlh-3* function (Figure7, Figure 8A). The proximal VCs express *plin-11*::*mCherry* at higher proportions than the distal VCs. This prompted us to ask whether the presence of *lin-11* transcriptional activity is dependent on a secondary pathway other than one that is mediated by *hlh-3*. Given that others have shown *lin-11* acts downstream of EGF, *lin-11* may be targeted by both a *hlh-3* dependent pathway and this secondary EGF-dependent pathway (Figure 8B; Zheng et al., 2013).

We propose that the reason *lin-11* transcriptional activity is observed in the proximal VCs of *hlh-3 (lf)* individuals is that EGF-dependent signaling is acting in parallel to *hlh-3*. It is known that the proximal VCs acquire this subclass-specific identity feature (*unc-4*) in a time-dependent manner, occurring after EGF signaling, after mid-L4 development (Zheng et al., 2013). Our analysis suggests the EGF signaling pathway promotes *lin-11* transcription too. This would explain why, in the absence of *hlh-3*, there is still expression of *lin-11* (Figure 7). Lastly, our findings that *hlh-3 (lf)* mutants also exhibit reduced *unc-4* transcriptional activity in the proximal VCs is a logical consequence of lower *lin-11* expression in the proximal VCs (Figure 4E, Figure 8A). Our model shows that two pathways affect the expression of *lin-11* and other VC identity genes (Figure 8).

### *hlh-3* may be a terminal selector of VC fate

*hlh-3* meets several criteria to be classified as a gene encoding a terminal selector, in the VCs First, it is expressed from the birth to the maturation of all VC features. Second, in its absence, all known VC class terminal identity features fail to be acquired. Lastly, it functions cell-autonomously. Since more than one terminal selector can function to regulate downstream effector genes, it is possible that another terminal selector may function with *hlh-3*. To confirm if *hlh-3* is a terminal selector, additional work will need to test for the direct regulation of VC identity genes by *hlh-3*.

## MATERIALS AND METHODS

### Strain maintenance

All strains were maintained at 22°C on nematode growth media using standard conditions (Brenner, 1974). Some strains were provided by the CGC, which is funded by NIH Office of Research Infrastructure Programs (P40 OD010440). *hlh-3 (tm1688)* was isolated by the National Bioresource Project of Japan. *cccIs1* was kindly shared by Dr. Jennifer Ross Wolf, *uIs45* was kindly shared by Dr. Martin Chalfie, *otIs456* was kindly shared by Dr. Oliver Hobert, and *otIs564* was kindly shared by Dr. Paschalis Kratsios. See Supplementary Table 1 for a complete list of strains used in this study.

### Construction of transgenic strains

The transgenic strain harboring *icIs270* was generated by the integration of *akEx31 [pglr-5::gfp + lin-15(+)]* using UV-TMP treatment followed by outcrossing (see below). The VC rescue array *icEx274 [VC::hlh-3cDNA::GFP; pmyo-2::mCherry]* was generated by co-injection of the constructs pCFJ90 *(pmyo-2::mCherry)* and pRD2 *(VC::hlh-3cDNA::GFP)* into the mutant strain harboring *hlh-3 (tm1688); otIs45* at 20 ng/microliter and 2 ng/microliter, respectively. pRD2 was generated by Dr. Ryan Doonan to address whether *hlh-3* could rescue the egg-laying defective phenotype in *hlh-3 (tm1688)* (Doonan, 2006). The pRD2 construct contains a VC specific promoter obtained from the vector pDM4, kindly provided by Dr. Michael Koelle driving expression of a *hlh-3* cDNA (Doonan, 2006).

### Integration of extrachromosomal arrays

The transgenic strain harboring *icIs270* was generated by exposing L4 hermaphrodites to UV-TMP (350microJoules x 100 on Stratagene UV Stratalinker; 0.03microgram/microliter TMP. Irradiated animals were placed onto seeded NGM plates and transferred the next day to fresh seeded NGM plates (3 Po/plate). These were followed to clone F1s (∼150) and subsequently to clone three F2s per F1.

### Construction of HLH-3::GFP CRISPR-Cas-9 engineered line [ic272]

Construction of the CRISPR line required modification of two plasmids: the single guide RNA or sgRNA plasmid, pDD162 (Addgene #47549), and the repair template plasmid, pDD282 (Addgene #66823) (Dickinson et al., 2015). The target sequence GCTATGATGATCACCAGAAG was selected using the CRISPR design tool on Flybase consisting of a high optimal quality score (96). The sgRNA was cloned into pDD162 to create pLP1. The 5’arm homology arm was designed as a gBlock containing a silent mutation at the PAM site to prevent Cas-9 off-targeting. The gBlock was PCR amplified with primers acgttgtaaaacgacggccagtcgccggca and CATCGATGCTCCTGAGGCTCCCGATGCTCC and cloned into pDD282. The 3’ homology arm was designed via PCR using the primers CGTGATTACAAGGATGACGATGACAAGAGATAATCTGTTAAGTTGTACC and ggaaacagctatgaccatgttatcgatttccaaggagctggtgcacaag. The PCR product was purified and cloned into pDD282 to create pLP2. The modified constructs pLP1and pLP2, as well as the co-injection plasmid pGH8 (Addgene #19359) were co-injected into an N2 strain: sg-RNA plasmid (pLP1) at 50ng/uL; *hlh-3* repair template plasmid (pLP2) at 10ng/uL, and pGH8 at 2.5ng/uL. Screening was carried out according to the published protocol (Dickinson et al., 2015).

### Microscopy

Animals were mounted on 3% agarose pads containing droplets of 10mM levamisole. Fluorescent images were acquired with AxioVision on Zeiss Axioskop 2 microscope. Following the collection of images, some conversions were made with FIJI version 2.0.0 (grayscale images were converted with Lookup tables: Red or Green) and processed into Adobe Illustrator for formatting. Fluorescent reporters were observed under confocal microscopy for the detection of a fluorescent protein signal (presence or absence) in transgenic lines. This study does not report quantification of intensity for any fluorescent reporter observed.

## ACKNOWLEDGMENTS

We would like to acknowledge Freddy Jacome, Basil Muhana, and Alex Obafemi for help in data acquisition; Dr. Suzanne McCutcheon for imaging equipment; and the National Science Foundation Bridge to the Doctorate Fellowship as well as the Department of Biological Sciences for support of LMP. We thank Martin Chalfie, Oliver Hobert, Paschalis Kratsios, and Jennifer Wolff for kindly sharing strains. We are also grateful to Kimberly Goodwin for their comments on the manuscript.

## SUPPLEMENTARY INFORMATION

**Supplementary Figure 1.**
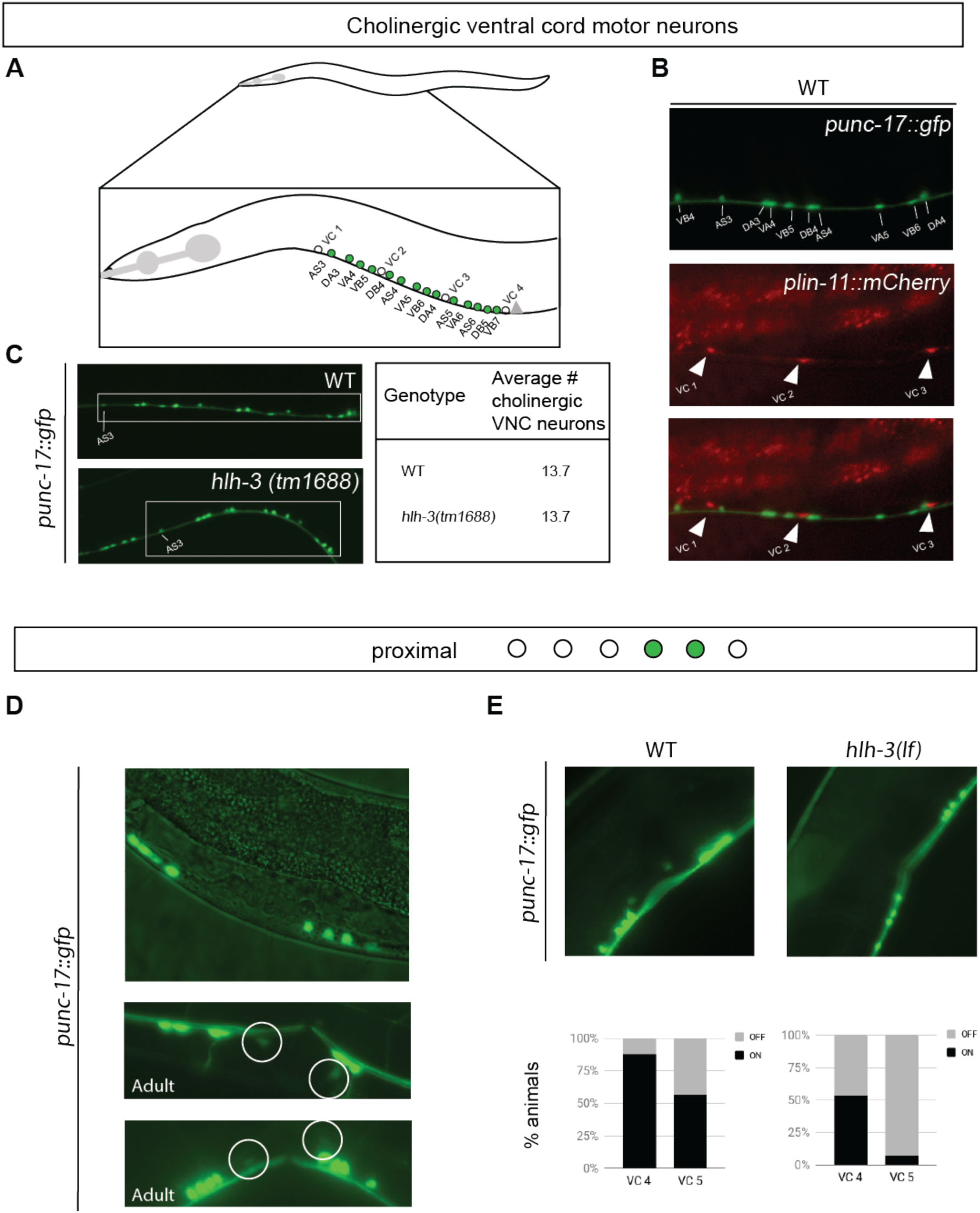
Cholinergic, sex-shared ventral cord motor neurons differentiate normally in *hlh-3lf*. **A:** Schematic of the number and position of cholinergic, sex-shared VNC neurons in the anterior body region, and between VC 1 and VC 4 (n = 14). **B:** An annotated image of adult WT hermaphrodite expressing *punc-17::gfp* in non-VC neurons (top panel), *plin-11::mCherry* in VCs (middle panel, filled arrowheads), and a merge of both images (bottom image). Anterior is left, ventral is down. **C:** Quantification of number of *punc-17::gfp* positive nuclei in the anterior region of the vulva in WT (n = 10) and *hlh-3 (lf)* (n = 10) hermaphrodites. Representative images are shown on the left. The average number of positive nuclei is reported on the right for each genotype. **D:** Representative images of L4 and adult WT hermaphrodites harboring the *punc-17::gfp* (*vsIs48*) reporter. There is no detectable expression in mid-L4 development (top panel), but the expression is detected in adults (middle and bottom panels). **E:** Quantification of reporter expression in proximal VCs of WT (n = 16) and *hlh-3 (lf)* (n = 15) in adulthood. On = detectable, Off = undetectable.

**Supplementary Figure 2.**
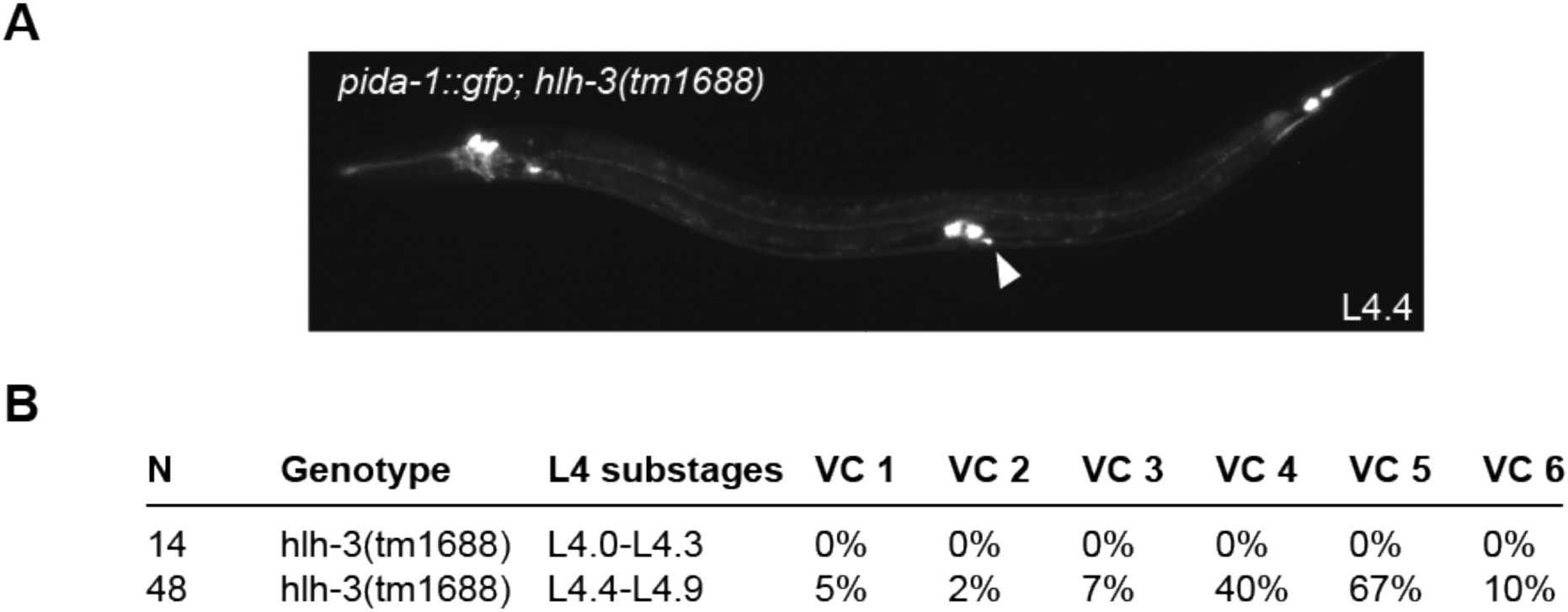
*hlh-3* acts prior to early larval L4 substages. **A:** Image of an early L4 *hlh-3 (lf)* hermaphrodite expressing *pida-1::gfp* only in VC 5 (white arrowhead). In WT individuals this reporter is detectable in all VCs (Figure 2) as well as the round-shaped bodies near the vulva, a pair of uv1 cells. Expression in uv1 cells is not affected in *hlh-3 (lf)* individuals **B:** Quantification analysis of *pida-1::gfp* detection in each VC of *hlh-3 (lf)* individuals during early L4 substages (L4.0-L4.3) or mid-late substages (L4.4-L4.9).

**Supplementary Figure 3.**
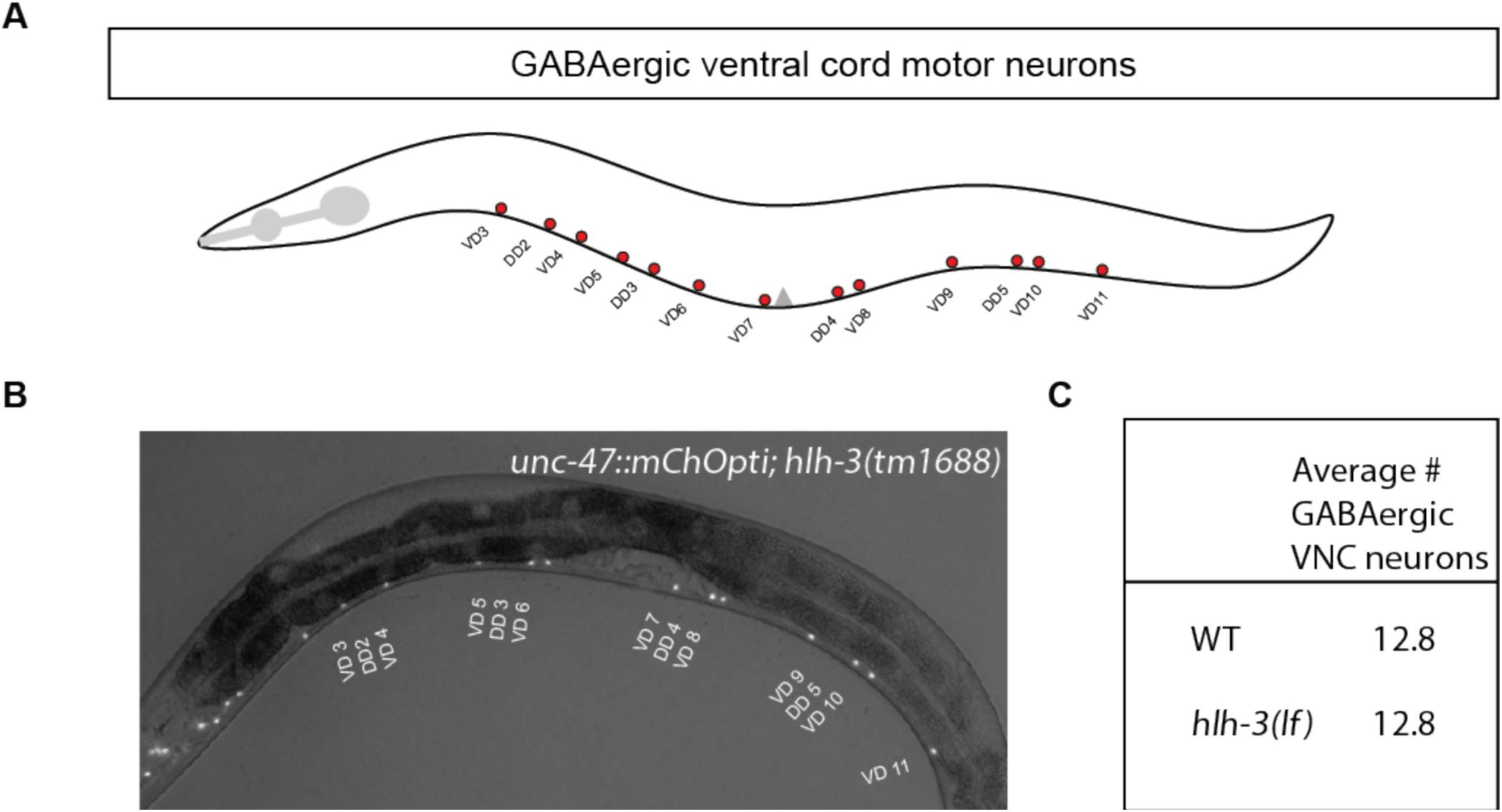
GABAergic, sex-shared, ventral cord motor neurons differentiate normally in *hlh-3 (lf)*. **A:** Illustration of the positions of the GABAergic VNC motor neurons scored (only VD 3 through VD 11 were scored, n = 13). **B:** Representative image of *unc-4*7 reporter expression (*otIs564 [unc-47fosmid::SL2::mChOpti::H2B; pha-1(+)]*) in a *hlh-3 (lf)* mutant individual in L4 development. The gene *unc-47* encodes a vesicular GABA transporter; it marks GABAergic neurons in the VNC. Both WT and *hlh-3 (lf)* individuals express the *unc-47* marker (WT not shown). **C:** Quantification of VNC neurons expressing *otIs564* reported as averages per genotype in one day old WT (n = 14) and *hlh-3 (lf)* (n = 14) hermaphrodites.

**Supplementary Figure 4.**
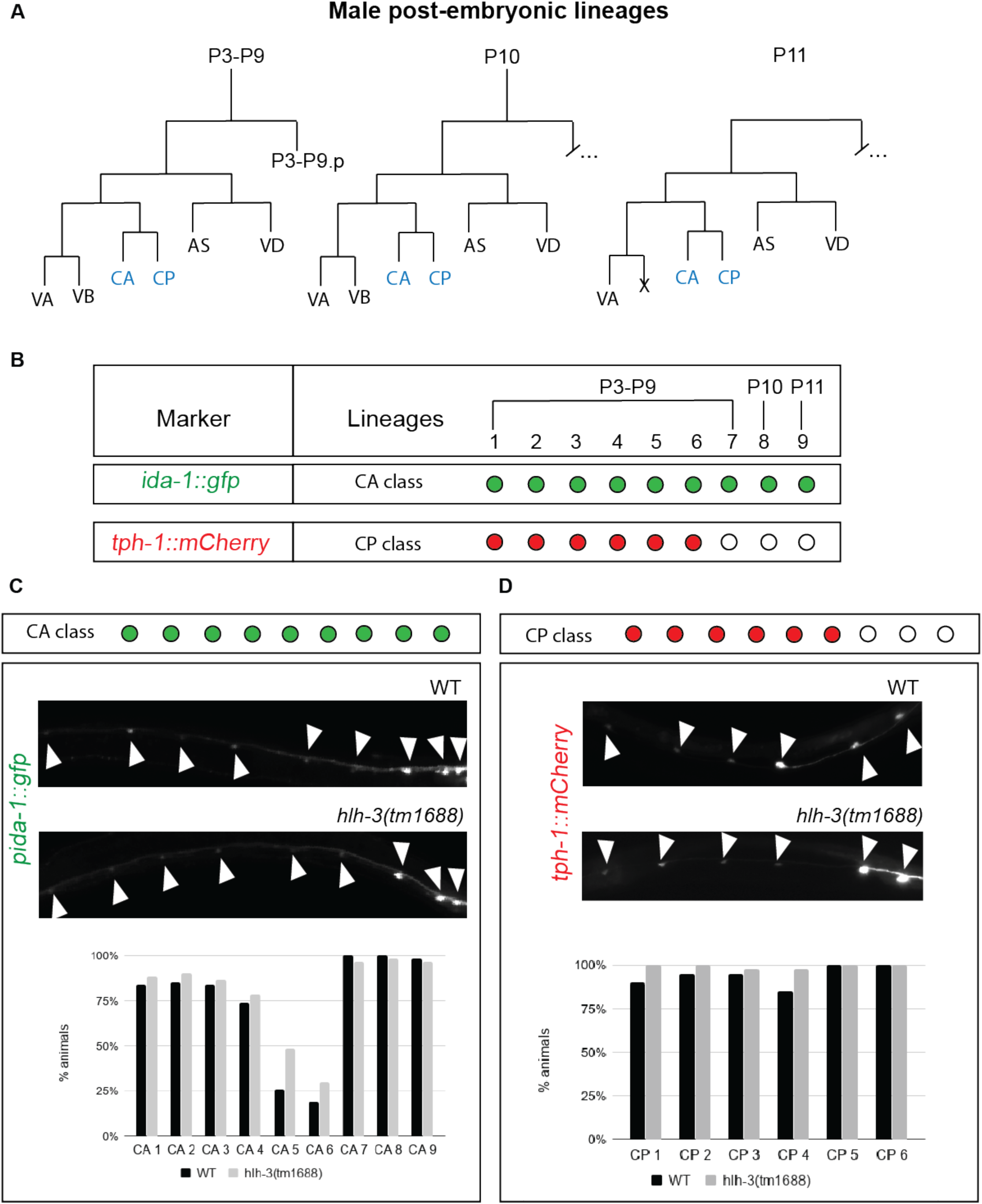
The differentiation of the male-specific ventral cord motor neurons derived from P cells is not affected by the absence of *hlh-3* function. **A:** Diagram of post-embryonic lineages in the ventral nerve cord that gives rise to CA and CP male-specific neurons. Notably, P2.a divisions give rise to CP0 but are not shown (adapted from Sulston et al., 1980). **B:** Summary of the expression pattern of *ida-1::gfp* and *tph-1::mCherry* in CAs and CPs, respectively (based on data from Kalis et al., 2014; Loer & Kenyon, 1993). **C:** Quantification of expression of *pida-1::gfp* in the adult male ventral cord of wild type and mutant individuals. Representative fluorescent images for each genotype (top). Graph reports the percent of animals with detectable expression in each cell of WT (n = 71) and *hlh-3 (lf)* (n = 61) males. **D:** Quantification of expression of *ptph-1::mCherry* expression in the adult male ventral cord of wild type and mutant individuals. Representative fluorescent images for each genotype (top). Graph reports the percent of animals with detectable expression in each cell of WT (n = 20) and *hlh-3 (lf)* (n = 41) males.

---

**Supplementary Table 1.**
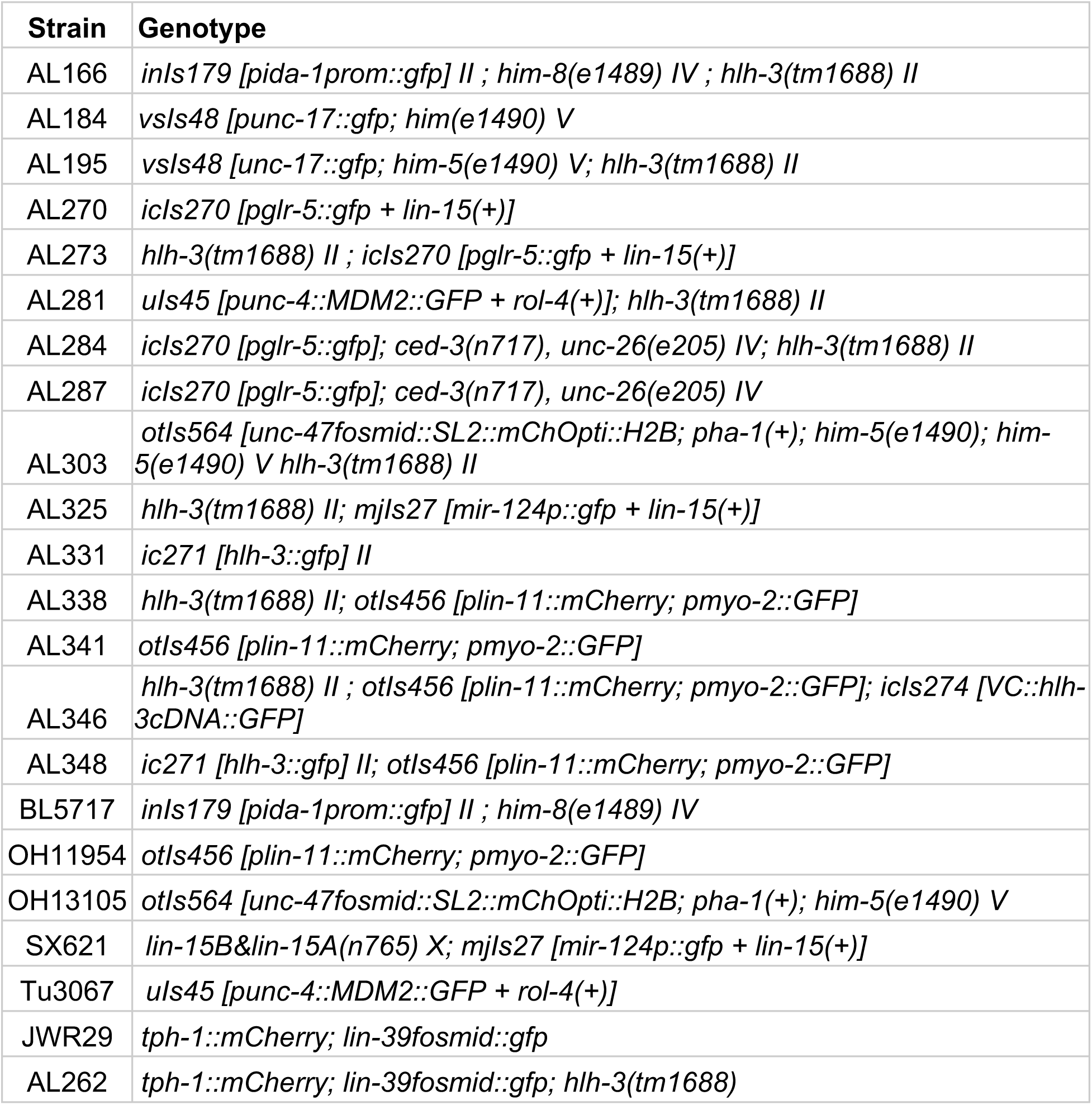
List of Strains

| Strain | Genotype |
| --- | --- |
| AL166 | *inls179 [pida-1prom::gfp] II ; him-8(e1489) IV ; hlh-3(tm1688) II* |
| AL184 | *vsIs48 [punc-17::gfp; him(e1490) V* |
| AL195 | *vsIs48 [unc-17::gfp; him-5(e1490) V; hlh-3(tm1688) II* |
| AL270 | *icls270 [pglr-5::gfp + lin-15(+)]* |
| AL273 | *hlh-3(tm1688) II ; icls270 [pglr-5::gfp + lin-15(+)]* |
| AL281 | *uls45 [punc-4::MDM2::GFP + rol-4(+)]; hlh-3(tm1688) II* |
| AL284 | *icls270 [pglr-5::gfp]; ced-3(n717), unc-26(e205) IV; hlh-3(tm1688) II* |
| AL287 | *icls270 [pglr-5::gfp]; ced-3(n717), unc-26(e205) IV* |
| AL303 | *otIs564 [unc-47fosmid::SL2::mChOpti::H2B; pha-1(+); him-5(e1490); him-5(e1490) V hlh-3(tm1688) II* |
| AL325 | *hlh-3(tm1688) II; mjIs27 [mir-124p::gfp + lin-15(+)]* |
| AL331 | *ic271 [hlh-3::gfp] II* |
| AL338 | *hlh-3(tm1688) II; otIs456 [plin-11::mCherry; pmyo-2::GFP]* |
| AL341 | *otIs456 [plin-11::mCherry; pmyo-2::GFP]* |
| AL346 | *hlh-3(tm1688) II ; otIs456 [plin-11::mCherry; pmyo-2::GFP]; icls274 [VC::hlh-3cDNA::GFP]* |
| AL348 | *ic271 [hlh-3::gfp] II; otIs456 [plin-11::mCherry; pmyo-2::GFP]* |
| BL5717 | *inls179 [pida-1prom::gfp] II ; him-8(e1489) IV* |
| OH11954 | *otIs456 [plin-11::mCherry; pmyo-2::GFP]* |
| OH13105 | *otIs564 [unc-47fosmid::SL2::mChOpti::H2B; pha-1(+); him-5(e1490) V* |
| SX621 | *lin-15B&lin-15A(n765) X; mjIs27 [mir-124p::gfp + lin-15(+)]* |
| Tu3067 | *uls45 [punc-4::MDM2::GFP + rol-4(+)]* |
| JWR29 | *tph-1::mCherry; lin-39fosmid::gfp* |
| AL262 | *tph-1::mCherry; lin-39fosmid::gfp; hlh-3(tm1688)* |

